# Learning retention mechanisms and evolutionary parameters of duplicate genes from their expression data

**DOI:** 10.1101/2020.06.19.162107

**Authors:** Michael DeGiorgio, Raquel Assis

## Abstract

Learning about the roles that duplicate genes play in the origins of novel phenotypes requires an understanding of how their functions evolve. To date, only one method—`CDROM`—has been developed with this goal in mind. In particular, CDROM employs gene expression distances as proxies for functional divergence, and then classifies the evolutionary mechanisms retaining duplicate genes from comparisons of these distances in a decision tree framework. However, `CDROM` does not account for stochastic shifts in gene expression or leverage advances in contemporary statistical learning for performing classification, nor is it capable of predicting the underlying parameters of duplicate gene evolution. Thus, here we develop `CLOUD`, a multi-layer neural network built upon a model of gene expression evolution that can both classify duplicate gene retention mechanisms and predict their underlying evolutionary parameters. We show that not only is the `CLOUD` classifier substantially more powerful and accurate than `CDROM`, but that it also yields accurate parameter predictions, enabling a better understanding of the specific forces driving the evolution and long-term retention of duplicate genes. Further, application of the `CLOUD` classifier and predictor to empirical data from *Drosophila* recapitulates many previous findings about gene duplication in this lineage, showing that new functions often emerge rapidly and asymmetrically in younger duplicate gene copies, and that functional divergence is driven by strong natural selection. Hence, `CLOUD` represents the best available method for classifying retention mechanisms and predicting evolutionary parameters of duplicate genes, thereby also highlighting the utility of incorporating sophisticated statistical learning techniques to address long-standing questions about evolution after gene duplication.

## Introduction

Gene duplication is a mutational process that creates copies of existing genes. Experimental studies in several diverse species have revealed that duplication occurs faster than all other types of spontaneous mutation [Lynch et al., 2008, Lipinski et al., 2011, Schrider et al., 2013, Keith et al., 2016, Konrad et al., 2018], thus serving as a major reservoir of genetic variation. Moreover, in contrast to other types of mutation, duplication generates redundancy, permitting the exploration of evolutionary space that may have been ancestrally forbidden [Ohno, 1970]. As a result, duplication has long been hypothesized to underlie the origins of novel phenotypes and complex biological systems [Ohno, 1970]. Indeed, surmounting evidence of widespread duplication and its contribution to adaptation and speciation in all three biological kingdoms [Zhang, 2003, Kondrashov, 2012] highlights its key role in evolution across the tree of life.

Yet, the evolutionary path from a pair of redundant gene copies to two functionally-independent genes remains unclear (Figure 1). Though redundancy can promote adaptation through a relaxation of selective constraint [Ohno, 1970], beneficial mutations are rare [Lynch and Force, 2000]. Hence, theory predicts that the most common outcome of duplication is nonfunctionalization, whereby one gene copy loses its function via an accumulation of deleterious mutations, leading to a reversion back to the ancestral singlecopy state [Lynch and Force, 2000]. As a result, four mechanisms have been proposed to explain how numerous duplicate genes bypass nonfunctionalization and are retained over millions of years of evolution [Ohno, 1970, Zhang, 2003, Force et al., 1999, Stoltzfus, 1999, He and Zhang, 2005, Rastogi and Liberles, 2005]. First, either benefits of increased gene dosage [Ohno, 1970] or recombination between gene copies [Zhang, 2003] may result in conservation, whereby both copies maintain the ancestral function. Second, beneficial mutations arising in one gene copy may lead to neofunctionalization, whereby this copy acquires a new function while the other maintains the ancestral function [Ohno, 1970]. Third, deleterious mutations targeting different functional domains of each gene copy may result in subfunctionalization, whereby each copy maintains a distinct subset of the ancestral function [Force et al., 1999, Stoltzfus, 1999]. Fourth, a combination of deleterious and beneficial mutations targeting different functional domains of each gene copy may lead to specialization, whereby each copy maintains a subset of the ancestral function and also acquires a new function [He and Zhang, 2005, Rastogi and Liberles, 2005].

**Figure 1:**
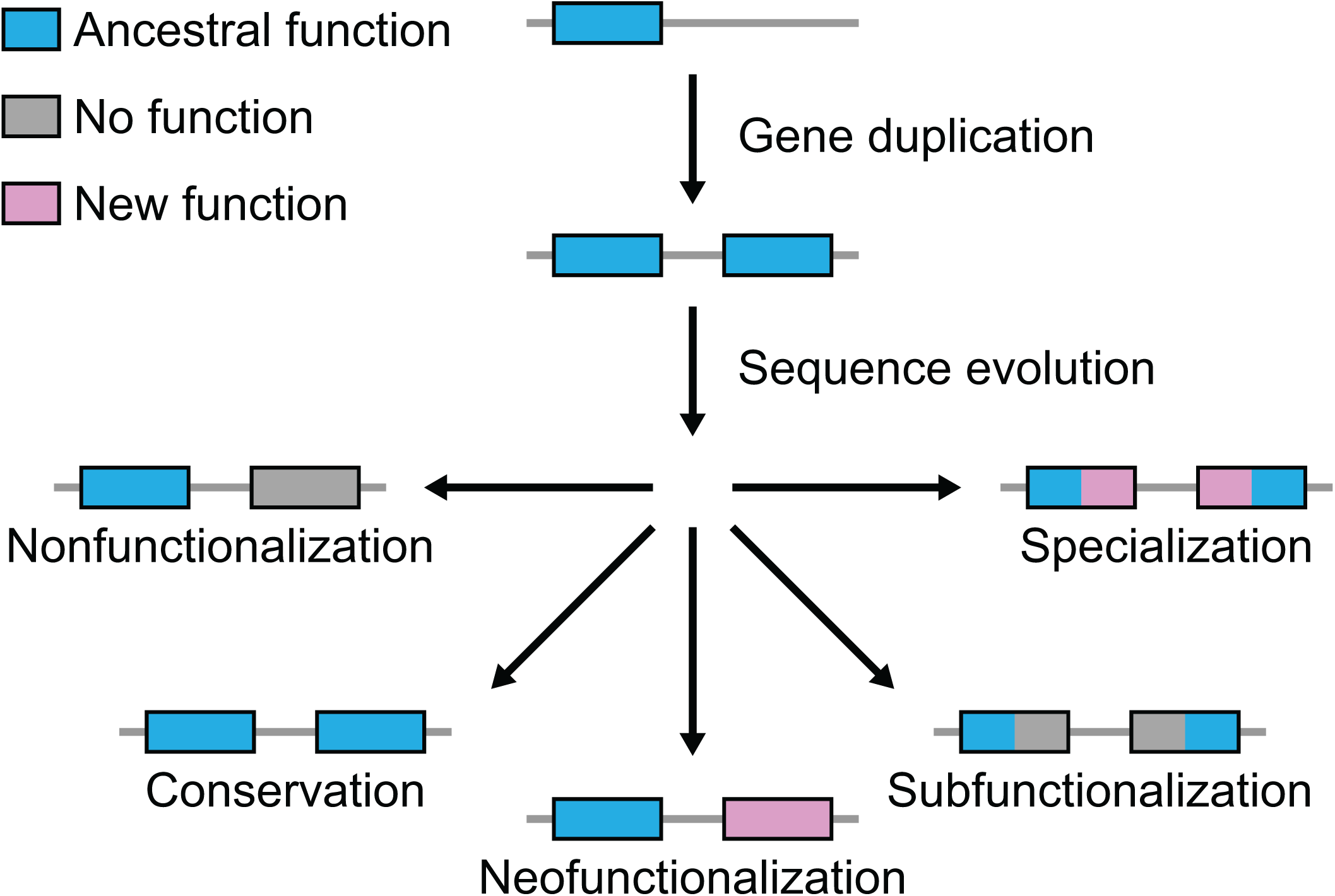
Hypothesized evolutionary trajectories of duplicate genes. Gene duplication results in two copies of an ancestral gene. Evolution may result in the loss of one functional copy by nonfunctionalization, or in the retention of two functional copies by either conservation, neofunctionalization, subfunctionalization, or specialization.

Due to their vastly different evolutionary forces and functional outcomes, differentiating among these four retention mechanisms is critical to understanding how gene duplication drives phenotypic innovation. In previous work, Assis and Bachtrog [2013] tackled this problem by designing a decision tree classification algorithm based on comparisons of differences between multi-tissue expression profiles of ancestral singlecopy genes and their derived duplicate gene copies. This approach [Assis and Bachtrog, 2013], which was later generalized to other types of input data and implemented in the R package `CDROM` [Perry and Assis, 2016], has been used to classify retention mechanisms of duplicate genes in *Drosophila* [Assis and Bachtrog, 2013], mammals [Assis and Bachtrog, 2015], honeybees [Chau and Goodisman, 2017], and grasses [Jiang and Assis, 2019]. Together, these studies have demonstrated that duplicate genes are frequently retained by neofunctionalization [Assis and Bachtrog, 2013, Assis, 2014, Assis and Bachtrog, 2015, Chau and Goodisman, 2017, Jiang and Assis, 2019], that the young “child” copy more often acquires a new function than the original “parent” copy [Assis and Bachtrog, 2013, Assis, 2014, Assis and Bachtrog, 2015, Jiang and Assis, 2019], and that new functions tend to be male-specific [Assis and Bachtrog, 2013, Assis, 2014, Assis and Bachtrog, 2015, Chau and Goodisman, 2017, Jiang and Assis, 2019]. Further, a follow-up study in *Drosophila* revealed natural selection to play important roles in both whether and how duplicate genes are retained over evolutionary time [Jiang and Assis, 2017].

However, there are two major shortcomings of the method implemented by `CDROM` [Assis and Bachtrog, 2013, Perry and Assis, 2016]. First, it does not account for stochastic shifts in gene expression that may occur as a result of phenotypic drift [Oleksiak et al., 2002, Khaitovich et al., 2004]. Second, it does not leverage the power provided by recent advances in statistical and machine learning [Hastie et al., 2009, Goodfellow et al., 2016]. With these limitations in mind, we developed `CLOUD` (CLassification using Ornstein-Uhlenbeck of Duplicates), a novel classification algorithm that employs simulated training data generated by Ornstein-Uhlenbeck (OU) processes, which can model gene expression evolution along phylogenetic trees [Hansen, 1997, Butler and King, 2004, Bedford and Hartl, 2008, Kalinka et al., 2010, Brawand et al., 2011, Perry et al., 2012, Rohfls et al., 2014, Rohfls and Nielsen, 2015]. In particular, because OU processes model Brownian motion with a pull toward an optimal state, they have a natural application to evolution, in which phenotypic drift is analogous to Brownian motion, natural selection to pull, and fittest phenotype to optimal state [Hansen, 1997, Butler and King, 2004].

Though OU processes have been used to model expression evolution of single-copy genes [Bedford and Hartl, 2008, Kalinka et al., 2010, Brawand et al., 2011, Perry et al., 2012, Rohfls et al., 2014, Rohfls and Nielsen, 2015], they have never been applied to the analogous problem after gene duplication. Thus, `CLOUD` adapts the OU framework to quantify expression evolution after gene duplication by modeling changes along a tree relating a pair of duplicate genes (parent and child copies) and their ancestral gene in a related sister species. Then, it utilizes the simulated output of these models to construct a multi-layer feedforward neural network for classifying duplicate genes as retained under conservation, neofunctionalization, subfunctionalization, or specialization. Moreover, this approach enables `CLOUD` to also predict parameters influencing the expression evolution of duplicate genes. Application of `CLOUD` to simulated data shows that it has high power to differentiate among classes, vastly outperforming `CDROM` for a wide range of parameter values, and also accurately predicts parameters shaping the expression evolution of retained duplicate genes. Further, application of `CLOUD` to empirical data from *Drosophila* [Assis and Bachtrog, 2013, Assis, 2019] recapitulates a majority of the classified duplicate gene retention mechanisms presented by Assis and Bachtrog [2013], as well as generates parameter predictions that match theoretical expectations of these retention mechanisms. CLOUD has been implemented as an R package, and is freely available at http://assisgroup.fau.edu/software.html. Its input data can include gene expression measured for a single condition or multiple conditions of varying types (*e.g.*, tissues, developmental stages, or disease states), making it applicable to a wide range of single- and multi-cellular biological systems.

## Results

In this section, we design our `CLOUD` classifier and predictor, evaluate its performance on simulated data, and apply it to an empirical dataset from *Drosophila*. First, we introduce our OU framework for modeling expression evolution after gene duplication, which forms the basis of the `CLOUD` classifier and predictor. Next, we formally define the multi-layer neural network architecture implemented by `CLOUD` for both classification and prediction tasks. We then employ simulations to evaluate the relative classification powers and accuracies of `CDROM` and `CLOUD` across a wide range of parameters, as well as in more targeted regions of the parameter space. We also use these simulations to probe its accuracy in predicting parameters driving gene expression evolution after duplication, specifically its ability to estimate optimal gene expression, selection strength, and phenotypic drift for each of the classified retention mechanisms. Last, we apply `CLOUD` to empirical data from *Drosophila* [Assis and Bachtrog, 2013, Assis, 2019] to classify retention mechanisms and predict underlying evolutionary parameters after gene duplication in this lineage.

### Modeling expression evolution after gene duplication as an OU process

To design a model of expression evolution after gene duplication, we consider a pair of related species, Species 1 and Species 2, whose lineages diverged from that of a common ancestor at time *T*_PCA_ (Figure 2A-C). Suppose that the common ancestor had a single-copy gene that underwent duplication, giving rise to a pair of duplicate genes at time *T*_PC_ in the lineage of Species 1 after its divergence from the lineage of Species 2. Of the pair of duplicate genes in Species 1, we designate the copy corresponding to the original single-copy gene in the ancestor and in Species 2 as the parent, and the new copy that is absent in both the ancestor and Species 2 as the child. Further, suppose that optimal expression states for the parent, child, and ancestral genes are given by *θ*_P_, *θ*_C_, and *θ*_A_, respectively. Likewise, the optimal expression state for the single-copy gene in the ancestor prior to the divergence of Species 1 and Species 2 is given as *θ*_PCA_, and for the single-copy gene in the lineage of Species 1 before duplication occurred as *θ*_PC_. Additionally, assume that *θ*_PC_ = *θ*_PCA_ = *θ*_A_. We then model expression along the tree relating the parent, child, and ancestral genes as changing randomly through phenotypic drift with strength *σ*^2^, and toward the optimal expression state through selection with strength *α*, according to an OU process.

**Figure 2:**
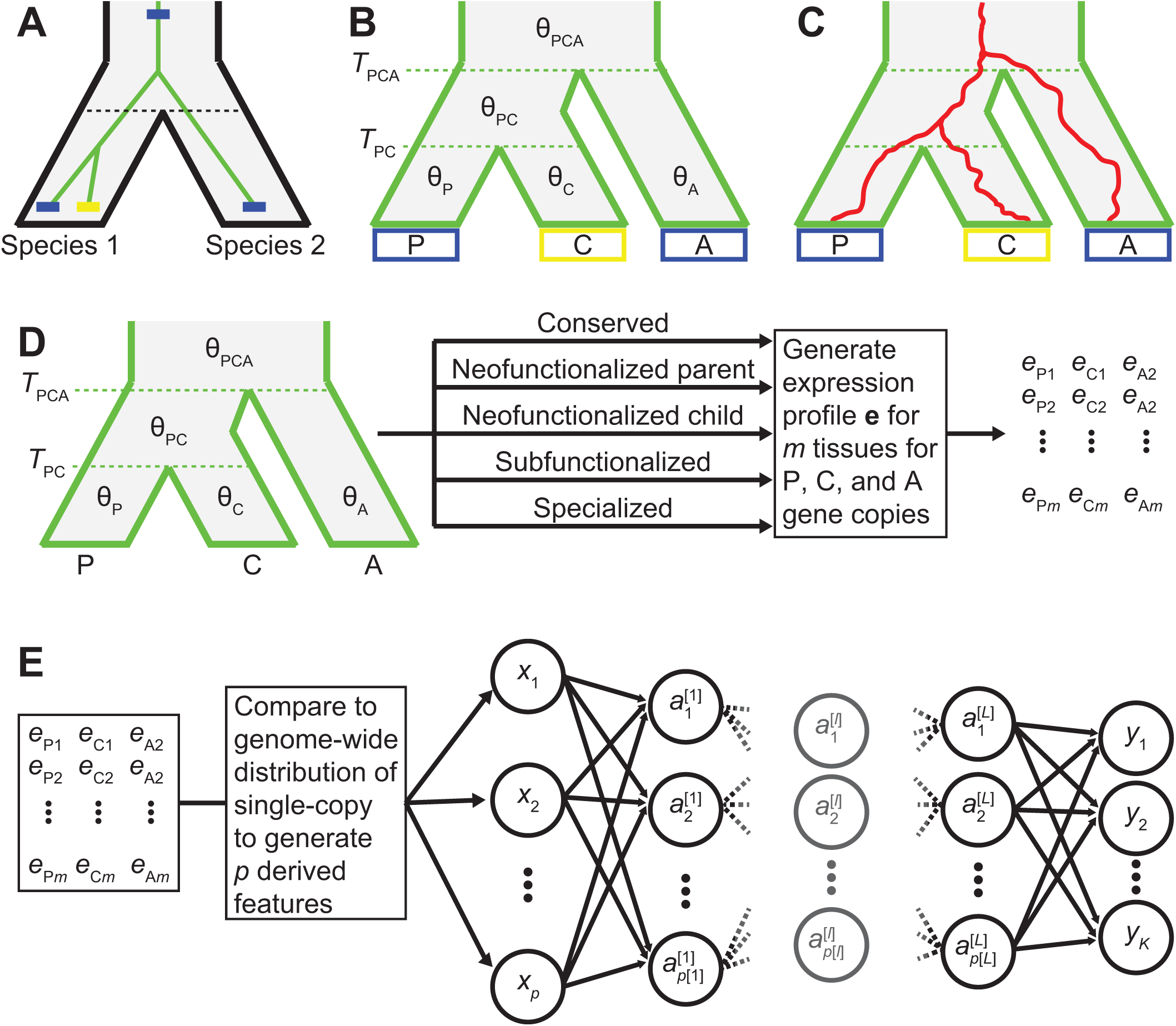
Modeling expression evolution after gene duplication as an OU process. (A) Relationships between two species (black phylogeny) and their genes (green phylogeny). After the two species diverged, a blue gene in Species 1 (parent) underwent a duplication event to create a yellow copy (child). (B) Relationships among the parent gene copy (P) in Species 1, child gene copy (C) in Species 1, and ancestral single-copy gene (A) in Species 2. The duplication event occurred at time *T*_PC_, and both copies split from the ancestral gene at time *T*_PCA_. Optimal expression states for the parent, child, and ancestral genes are given by *θ*_P_, *θ*_C_, and *θ*_A_, respectively. The internal branch and the branch above the root have optimal expression states *θ*_PC_ and *θ*_PCA_, respectively. (C) Cartoon depicting expression profile changes (red lines) along the gene tree. Expression profiles change randomly through phenotypic drift with strength *σ*^2^, and toward the optimal expression state through selection with strength *α*. (D) Illustration of how we simulate multi-tissue expression vectors for parent, child, and ancestral genes. (E) Schematic of our feed-forward neural network architecture, which takes in *p* input units with values *x*_1_, *x*_2_, …, *x*_*p*_, has *K* output units with values *y*_1_, *y*_2_, …, *y*_*K*_, and has *L* hidden layers, where the number of units in layer *𝓁* is *p*[*𝓁*] and the value of unit *k* in layer *𝓁* is the activation 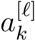.

In each tissue, gene expression **e** = (*e*_P_, *e*_C_, *e*_A_) ∈ ℝ^3^ is therefore distributed as a multivariate normal (MVN) normal distribution with mean

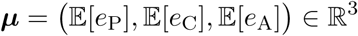

and covariance matrix

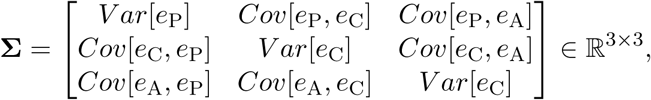

*i.e.*, **e** ∼ MVN(***µ*, Σ**). Following Brawand et al. [2011], we have that

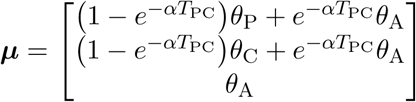

and

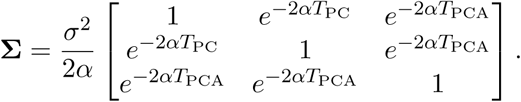

Here we assume that expression is independent across tissues. However, this approach can also be extended to account for the inter-tissue expression covariance structure using established approaches [Revell and Harmon, 2008, Revell and Collar, 2009, Clavel et al., 2015].

### Neural network architecture for the CLOUD classifier and predictor

We denote the set of all genes with two copies in one species and one copy in the other as duplicate genes *𝒟*, and the set of all genes with one copy in both species as single-copy genes *𝒢*. Let

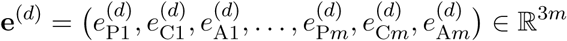

be the input expression vector for duplicate gene *d* ∈ *𝒟* across *m* tissues, where 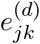 is the expression level for copy *j* ∈ {P, C, A} of duplicate gene *d* in tissue *k* ∈ {1, 2, …, *m*}. Similarly, let

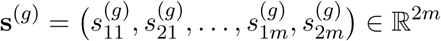

be the expression vector for single-copy gene *g* ∈ *𝒢* across *m* tissues, where 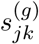 is the expression level for species *j* ∈ {1, 2} of single-copy gene *g* in tissue *k*.

We transform and compare the expression vector **e**^(*d*)^ of each duplicate gene *d* ∈ *𝒟* to the expression vector **s**^(*g*)^ of each single-copy gene *g* ∈ *𝒢* to obtain the feature vector

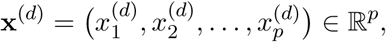

which we use as input to a dense feed-forward neural network. Following Assis and Bachtrog [2013], we compare multi-tissue expression differences between duplicate genes *𝒟* to the distribution of multi-tissue expression differences between single-copy genes *𝒢*. Specifically, we generate the set of *p* = 4*m* + 84 derived features listed in Table 1, many of which involve comparisons to distributions of values for single-copy genes *𝒢*. To generate these distributions, we compute the Euclidean distance and Pearson correlation coefficient between the multi-tissue expression vectors of Species 1 and Species 2 for each single-copy gene *g* ∈ *𝒢*. Based on these values, we derive the sets of all Euclidean distances dist(*𝒢*) and Pearson correlation coefficients cor(*𝒢*). We utilize features based on both dist(*𝒢*) and cor(*𝒢*) so that we can evaluate not only differences among values, but also among their shapes [Hastie et al., 2009].

**Table 1:** Set of p = 4m + 84 derived features used as input to CLOUD

| Feature number | Feature value |
| --- | --- |
| 1 | $t_{PC} = T_{PC}/T_{PCA}$ |
| 2 to $3m + 1$ | $\mathbf{e}^{(d)} = (e_{P1}^{(d)}, e_{C1}^{(d)}, e_{A1}^{(d)}, \dots, e_{Pm}^{(d)}, e_{Cm}^{(d)}, e_{Am}^{(d)})$ |
| $3m + 2$ to $4m + 1$ | $e_{P1}^{(d)} + e_{C1}^{(d)}, \dots, e_{Pm}^{(d)} + e_{Cm}^{(d)}$ |
| $4m + 2$ | $\text{dist}(P, C) = \sqrt{(e_{P1}^{(d)} - e_{C1}^{(d)})^2 + \dots + (e_{Pm}^{(d)} - e_{Cm}^{(d)})^2}$ |
| $4m + 3$ | $\text{dist}(P, A) = \sqrt{(e_{P1}^{(d)} - e_{A1}^{(d)})^2 + \dots + (e_{Pm}^{(d)} - e_{Am}^{(d)})^2}$ |
| $4m + 4$ | $\text{dist}(C, A) = \sqrt{(e_{C1}^{(d)} - e_{A1}^{(d)})^2 + \dots + (e_{Cm}^{(d)} - e_{Am}^{(d)})^2}$ |
| $4m + 5$ | $\text{dist}(PC, A) = \sqrt{(e_{P1}^{(d)} + e_{C1}^{(d)} - e_{A1}^{(d)})^2 + \dots + (e_{Pm}^{(d)} + e_{Cm}^{(d)} - e_{Am}^{(d)})^2}$ |
| $4m + 6$ | $\text{branch}(P) = [\text{dist}(P, C) + \text{dist}(P, A) - \text{dist}(C, A)]/2$ |
| $4m + 7$ | $\text{branch}(C) = [\text{dist}(P, C) + \text{dist}(C, A) - \text{dist}(P, A)]/2$ |
| $4m + 8$ | $\text{branch}(A) = [\text{dist}(P, A) + \text{dist}(C, A) - \text{dist}(P, C)]/2$ |
| $4m + 9$ | rank of $\text{dist}(P, C)$ among $\text{dist}(\mathcal{G})$ |
| $4m + 10$ | rank of $\text{dist}(P, A)$ among $\text{dist}(\mathcal{G})$ |
| $4m + 11$ | rank of $\text{dist}(C, A)$ among $\text{dist}(\mathcal{G})$ |
| $4m + 12$ | rank of $\text{dist}(PC, A)$ among $\text{dist}(\mathcal{G})$ |
| $4m + 13$ to $4m + 20$ | $k$th moment of $[\text{dist}(P, C) - \text{dist}(\mathcal{G})]/\max\{\text{dist}(\mathcal{G})\}$ for $k = 1, 2, \dots, 8$ |
| $4m + 21$ to $4m + 28$ | $k$th moment of $[\text{dist}(P, A) - \text{dist}(\mathcal{G})]/\max\{\text{dist}(\mathcal{G})\}$ for $k = 1, 2, \dots, 8$ |
| $4m + 29$ to $4m + 36$ | $k$th moment of $[\text{dist}(C, A) - \text{dist}(\mathcal{G})]/\max\{\text{dist}(\mathcal{G})\}$ for $k = 1, 2, \dots, 8$ |
| $4m + 37$ to $4m + 44$ | $k$th moment of $[\text{dist}(PC, A) - \text{dist}(\mathcal{G})]/\max\{\text{dist}(\mathcal{G})\}$ for $k = 1, 2, \dots, 8$ |
| $4m + 45$ | $\text{cor}(P, C) = \text{PearsonCorrelation}(e_{P1}^{(d)}, \dots, e_{Pm}^{(d)}; e_{C1}^{(d)}, \dots, e_{Cm}^{(d)})$ |
| $4m + 46$ | $\text{cor}(P, A) = \text{PearsonCorrelation}(e_{P1}^{(d)}, \dots, e_{Pm}^{(d)}; e_{A1}^{(d)}, \dots, e_{Am}^{(d)})$ |
| $4m + 47$ | $\text{cor}(C, A) = \text{PearsonCorrelation}(e_{C1}^{(d)}, \dots, e_{Cm}^{(d)}; e_{A1}^{(d)}, \dots, e_{Am}^{(d)})$ |
| $4m + 48$ | $\text{cor}(PC, A) = \text{PearsonCorrelation}(e_{P1}^{(d)} + e_{C1}^{(d)}, \dots, e_{Pm}^{(d)} + e_{Cm}^{(d)}; e_{A1}^{(d)}, \dots, e_{Am}^{(d)})$ |
| $4m + 49$ | rank of $\text{cor}(P, C)$ among $\text{cor}(\mathcal{G})$ |
| $4m + 50$ | rank of $\text{cor}(P, A)$ among $\text{cor}(\mathcal{G})$ |
| $4m + 51$ | rank of $\text{cor}(C, A)$ among $\text{cor}(\mathcal{G})$ |
| $4m + 52$ | rank of $\text{cor}(PC, A)$ among $\text{cor}(\mathcal{G})$ |
| $4m + 53$ to $4m + 60$ | $k$th moment of $\text{cor}(P, C) - \text{cor}(\mathcal{G})$ for $k = 1, 2, \dots, 8$ |
| $4m + 61$ to $4m + 68$ | $k$th moment of $\text{cor}(P, A) - \text{cor}(\mathcal{G})$ for $k = 1, 2, \dots, 8$ |
| $4m + 69$ to $4m + 76$ | $k$th moment of $\text{cor}(C, A) - \text{cor}(\mathcal{G})$ for $k = 1, 2, \dots, 8$ |
| $4m + 77$ to $4m + 84$ | $k$th moment of $\text{cor}(PC, A) - \text{cor}(\mathcal{G})$ for $k = 1, 2, \dots, 8$ |

Given the input feature vector **x**^(*d*)^, we seek to predict the output vector

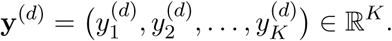

When performing classification of duplicate gene retention mechanisms, **y**^(*d*)^ is the vector of *K* = 5 class probabilities, corresponding to class labels “Conserved” for conservation, “Neofunctionalized parent” for neofunctionalization in which the parent copy acquires a new function, “Neofunctionalized child” for neofunctionalization in which the child copy acquires a new function, “Subfunctionalized” for subfunctionalization, and “Specialized” for specialization. In contrast, when predicting evolutionary parameters of duplicate genes, **y**^(*d*)^ is the vector of *K* = 5*m* parameter predictions in each of the *m* tissues, where in each tissue we obtain parameter estimates for *θ*_P_, *θ*_C_, *θ*_A_, *σ*^2^, and *α*.

We consider a dense feed-forward neural network with *L* ∈ {0, 1, 2, 3} hidden layers. The first hidden layer has *p*[1] = 256 hidden units, and hidden layer 𝓁 ∈ {1, 2, …, *L*} has *p*[𝓁] = 256*/*2^−1^ hidden units, such that each hidden layer has half the number of hidden units as the previous hidden layer. For the purposes of condensing notation, we also consider the input layer as hidden layer zero, such that *p*[0] = *p* = 4*m* + 84 is the number of input features, and we consider the output layer as hidden layer *L*+1, such that *p*[*L*+1] = *K*.

We define the values at unit *k* ∈ {1, 2, …, *p*[𝓁]} of hidden layer [𝓁 ∈ {0, 1, 2, …, *L*} for duplicate gene *d* ∈ *𝒟* by its activation 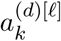. Because hidden layer zero is the input layer and hidden layer *L* + 1 is the output layer, then

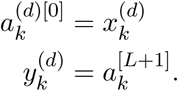

For hidden layer 𝓁 ∈ {1, 2, …, *L*}, we define the activation for unit *k* as a linear combination of the activations from the previous hidden layers, followed by a non-linear transformation [Goodfellow et al., 2016]. Here we choose the rectified linear unit [ReLU; Goodfellow et al., 2016] function defined as ReLU(*x*) = max(0, *x*), such that the activation for unit *k* in hidden layer 𝓁 of duplicate gene *d* is

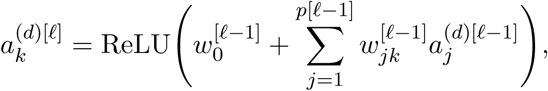

where 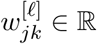; is the weight (parameter) from unit *j* in layer 𝓁 to unit *k* in layer 𝓁 + 1, and where 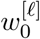 is the bias for layer 𝓁[Goodfellow et al., 2016]. The output layer takes inputs from layer *L*, and has a different form depending on whether we consider the classification or the prediction problem. For the classification problem, we employ the softmax activation function [Goodfellow et al., 2016], such that the output for class *k* ∈ {1, 2, …, *K*} of duplicate gene *d* is the probability

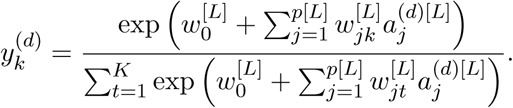

For the prediction problem, we instead use the linear activation function [Goodfellow et al., 2016], such that the output for parameter prediction *k* ∈ {1, 2, …, 5*m*} of duplicate gene *d* is

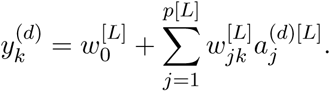

This neural network was implemented in R [R Core Team, 2013], using Keras [Chollet et al., 2017] with a TensorFlow backend [Abadi et al., 2015]. A schematic of the neural network architecture is provided in Figure 2E. Note that when *L* = 0, the neural network simplifies to a multinomial regression model [Hastie et al., 2009] for the classification problem, and to a linear regression model [Hastie et al., 2009] for the prediction problem.

### Classification power and accuracy of CLOUD relative to CDROM

To evaluate the classification power and accuracy of our multi-layer neural network classifier `CLOUD`, we trained and tested it on independent datasets simulated under each class of duplicate gene retention mechanisms (see *Methods*). We assumed two hidden layers when training and testing `CLOUD`, as this resulted in the best cross-validation performance (see *Methods*). Our training set consisted of 50,000 observations, of which 10,000 were simulated under each class. We trained `CLOUD` on these data, and explored evolutionary parameters drawn on a logarithmic scale across many orders of magnitude. Specifically, we independently drew parameters *θ*_P_, *θ*_C_, *θ*_A_ ∈ [10^−4^, 10^4^], *α* ∈ [1, 10^3^], and *σ*^2^ ∈ [10^−2^, 10^3^] across six tissues, for a total of 30 random parameters per simulated replicate. We then tested `CLOUD` on a separate set of 5,000 observations, of which 1,000 were simulated under each class, with evolutionary parameters drawn from the same broad space as that of the training set. For comparison, we also applied the existing classifier `CDROM` [Assis and Bachtrog, 2013, Perry and Assis, 2016] to the same simulated test set (see *Methods*).

Analysis of the resulting classifications reveals that `CLOUD` generally has substantially higher power (Figure 3A) and accuracy (Figure 3B) than `CDROM`. Specifically, across the wide range of test parameter values explored, `CLOUD` achieved an accuracy of 80.18%, whereas `CDROM` only reached 68.76% accuracy (Figure 3B). This represents a boost in over 16% accuracy of `CLOUD` above the baseline of the only previously available classifier CDROM. In addition to increased overall accuracy, `CLOUD` yields similar accuracies across classes, illustrated by narrow ranges of correct classification rates (between 77.1 and 84.2%; diagonal cells of Figure 3B) and mis-classification rates (between 1.3 and 9.2%; non-diagonal cells of Figure 3B). In contrast, `CDROM` demonstrates a much higher correct classification rate for the “Specialized” class (96.9%) than for other classes (between 45.7 to 67.9%; Figure 3B), and a higher mis-classification rate toward the “Specialized” class (between 28.7 to 30.6%; Figure 3B). Moreover, `CDROM` experiences additional issues when classifying true “Subfunctionalized” observations, with an 18.3% mis-classification rate toward the “Conserved” class (Figure 3B).

**Figure 3:**
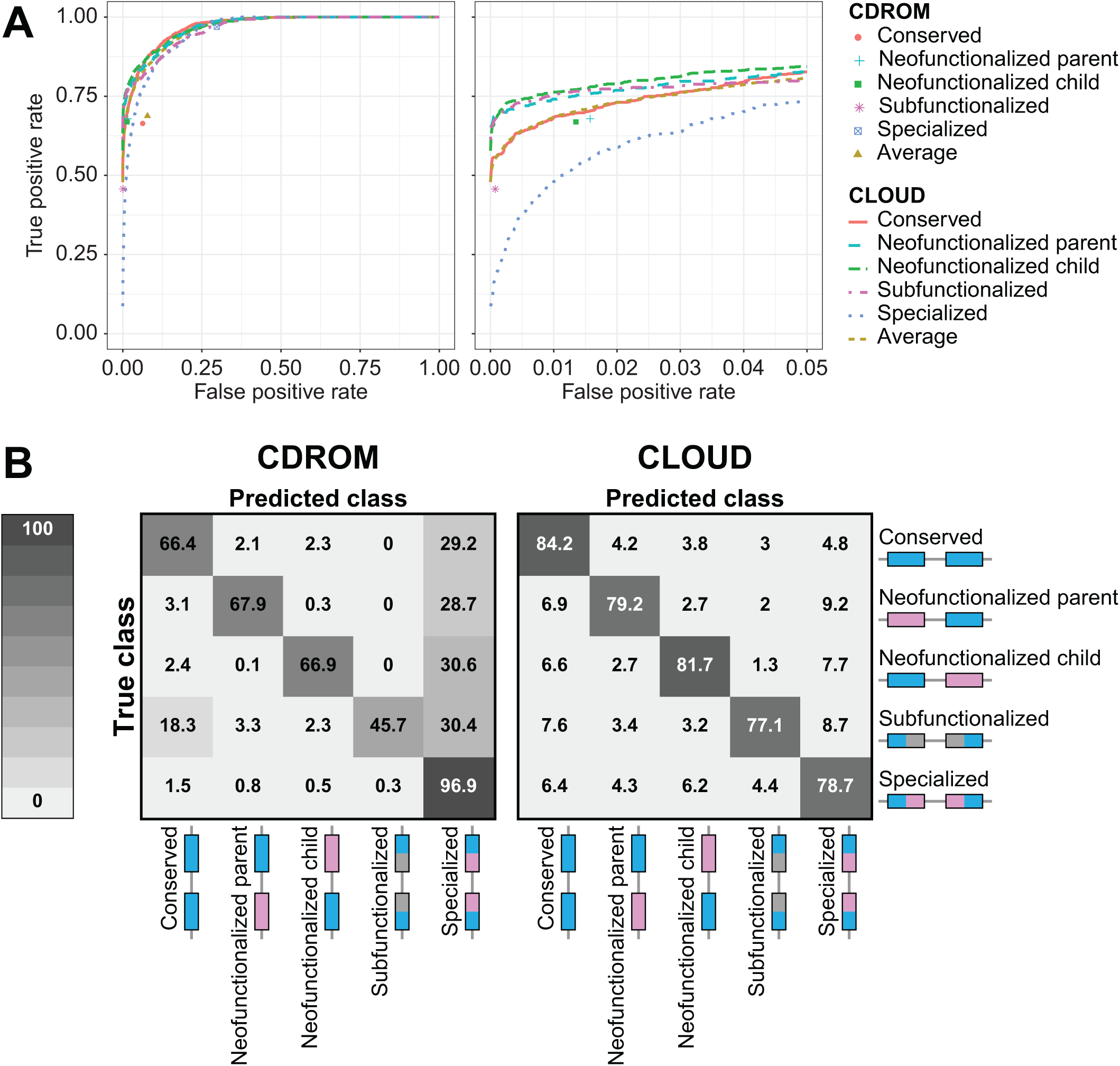
Classification results for `CDROM` and `CLOUD` with *L* = 2 hidden layers applied to data simulated under parameters *α* [1, 10^3^] and *σ*^2^ [10^−2^, 10^3^]. (A) Receiver operating characteristic curves across the full range of false positive rates (left) and truncated at a false positive rate of 5% (right). Because `CDROM` is a decision tree classifier, its true positive and false positive rates are plotted as points. (B) Confusion matrices depicting the classification rates of each of the five duplicate gene retention classes for `CDROM` (left) and `CLOUD` (right).

In addition, `CLOUD` is much more conservative than `CDROM` for pairs of *α* and *σ*^2^ values that are difficult to classify (Figures S3-S6). For example, both methods typically have higher power (Figures S5 and S6) and accuracy (Figures S3 and S4) when either selection is strong (large *α*) or random phenotypic drift is weak (small *σ*^2^). In contrast, when selection is weak (small *α*) and phenotypic drift is strong (large *σ*^2^), then classification is more difficult for both methods. However, in these cases, `CLOUD` tends to choose classes at similar rates, whereas `CDROM` is overconfident and chooses the “Specialized” class regardless of the true class (compare Figures S3 and S4). Therefore, `CLOUD` not only demonstrates uniformly higher power and accuracy than `CDROM` across a wide array of evolutionary settings, but is also unbiased unlike `CDROM`.

### Parameter prediction accuracy of CLOUD

In addition to its vastly improved classification performance relative to `CDROM`, a unique attribute of `CLOUD` is its ability to learn parameters underlying the expression evolution of duplicate genes. Thus, we next assessed the accuracy of the `CLOUD` predictor by training and testing it on the same independent simulated datasets that we employed for training and testing the `CLOUD` classifier. In particular, we trained `CLOUD` (again assuming two hidden layers) to make predictions for each of the five parameters (*θ*_P_, *θ*_C_, *θ*_A_, *α*, and *σ*^2^) in six tissues (total of 30 parameters) from the training set, and then applied it to make predictions for these parameters from the test set (see *Methods*).

To investigate prediction accuracy, we examined the distributions of mean parameter prediction errors across the six tissues (Figure 4). In general, all parameter estimates appear unbiased, with mean prediction errors centered on zero. Moreover, estimates of optimal expression states (*θ*_P_, *θ*_C_, and *θ*_A_) are more precise than those of selection strength (*α*), which are more precise than those of phenotypic drift (*σ*^2^). Further, parameter predictions for the “Specialized” class are less precise than those for other classes, likely due to the additional degrees of freedom in estimating parameters for this class. In particular, for the “Specialized” class, all optimal expression values are unconstrained (Table 2), whereas at least two of the three optimal expression states are constrained to be identical in the “Conserved” and “Neofunctionalized” classes, and *θ*_P_ and *θ*_C_ are constrained to sum to *θ*_A_ for the “Subfunctionalized” class (Table 2).

**Table 2:** Optimal expression states under OU processes used to simulate the five classes of duplicate gene retention mechanisms

| Class | Optimal expression state at a given tissue |
| --- | --- |
| Conserved | $\theta_P = \theta_C = \theta_A = \theta$ |
| Neofunctionalized parent | $\theta_C = \theta_A = \theta$ and $\theta_P \neq \theta$ |
| Neofunctionalized child | $\theta_P = \theta_A = \theta$ and $\theta_C \neq \theta$ |
| Subfunctionalized | $\theta_A = \theta$, $\theta_P \neq \theta$, $\theta_C \neq \theta$, and $\theta_P + \theta_C = \theta$ |
| Specialized | $\theta_A = \theta$, $\theta_P \neq \theta$, $\theta_C \neq \theta$, and $\theta_P + \theta_C \neq \theta$ |

**Figure 4:**
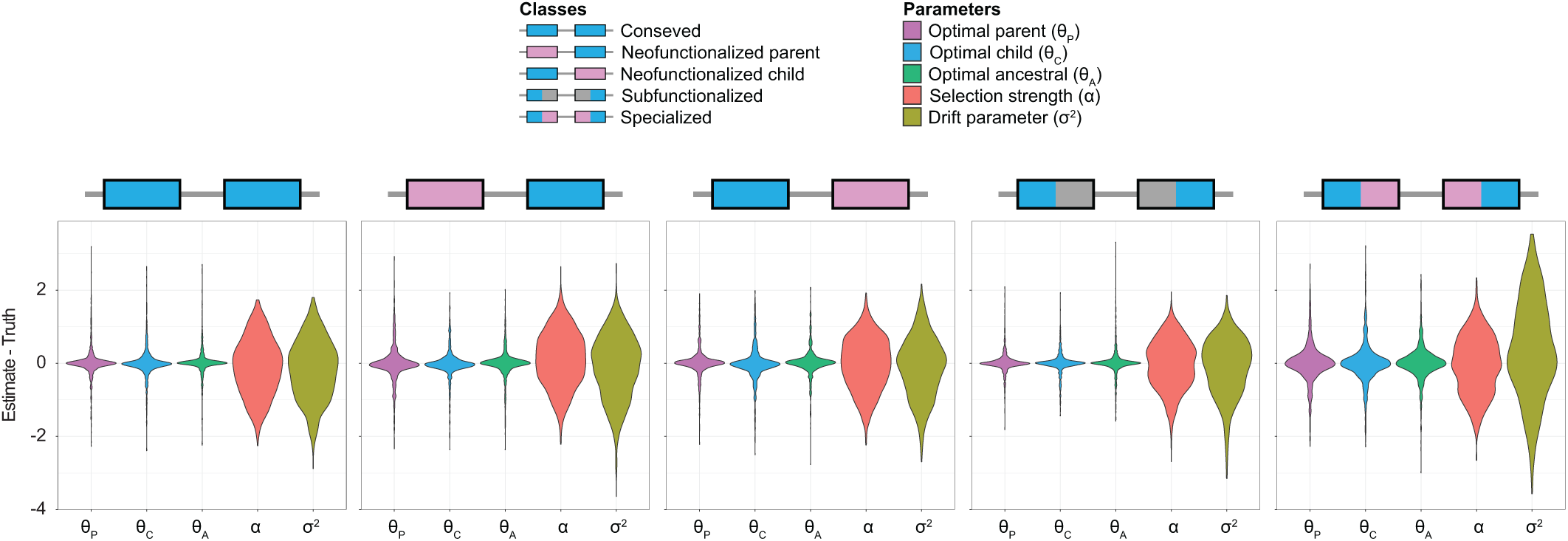
Prediction results for application of `CLOUD` with *L* = 2 hidden layers to data simulated under parameters *α* [1, 10^3^] and *σ*^2^ [10^−2^, 10^3^] for each of the five classes of duplicate gene retention mechanisms. Violin plots display distributions of parameter prediction errors averaged across the *m* = 6 tissues for each simulated test set.

As with classification, confidence in parameter predictions made by `CLOUD` also vary with *α* and *σ*^2^ (Figure S7). Though precision in estimation tends to be highest when selection is strong (large *α*) or phenotypic drift is weak (small *σ*^2^), it decreases as selection becomes weaker (smaller *α*) or phenotypic drift becomes stronger (larger *σ*^2^). Further, as with our general results across a wide parameter space (Figure 4), estimates of optimal expression states (*θ*_P_, *θ*_C_, and *θ*_A_) appear to be unbiased even in narrow regions of the space, with mean prediction errors centered on zero. In contrast, estimates of *α* and *σ*^2^ are biased for some pairs of values. Specifically, estimates of *α* and *σ*^2^ are both upwardly biased for weak selection (small *α*) with weak phenotypic drift (small *σ*^2^), and downwardly biased for strong selection (large *α*) with strong phenotypic drift (large *σ*^2^).

### Application of CLOUD to empirical data from *Drosophila*

Our simulation experiments highlight the exceptional classification performance of `CLOUD` relative to CDROM, as well as the unique ability of `CLOUD` to predict parameters underlying the evolution of duplicate genes. Hence, we next sought to use `CLOUD` to classify retention mechanisms and predict parameters of 208 duplicate genes in *Drosophila* [Assis and Bachtrog, 2013] from their expression data in six tissues [Assis, 2019]. Specifically, we first used PhyML [Guindon et al., 2010] to estimate a gene tree relating each parent, child, and ancestral gene in this dataset of duplicate genes [Assis and Bachtrog, 2013] (see *Methods*). Next, as in our simulation studies, we trained `CLOUD` (assuming two hidden layers) on a large balanced simulated training set of 50,000 observations (10,000 from each of five classes), with evolutionary parameters *θ*_P_, *θ*_C_, *θ*_A_ ∈ [−4, 4], log_10_(*α*) ∈ [0, 3], and log_10_(*σ*^2^) ∈ [−2, 3] drawn independently across six tissues, for a total of 30 random parameters per simulated training observation (see *Methods*). We tailored `CLOUD` to this dataset of duplicate genes [Assis and Bachtrog, 2013] by generating *p* = 108 input features (Table 1) from comparisons to the empirical distribution of single-copy genes identified in this lineage [Assis and Bachtrog, 2013] (see *Methods*). Then, we used `CLOUD` to classify retention mechanisms and predict parameters of the 208 *Drosophila* duplicate genes [Assis and Bachtrog, 2013] from these features.

Analysis of the resulting classifications reveals that the predominant mechanism of duplicate gene retention in *Drosophila* is neofunctionalization in which the child copy acquires a new function (61.43%; Figure 5), mirroring the findings of Assis and Bachtrog [2013]. Moreover, classifications of `CLOUD` are generally concordant with those of `CDROM` (59.29%), with three key differences. In particular, of the 167 duplicates classified as “Neofunctionalized child” by CDROM, 16 are classified as “Conserved” by `CLOUD`. In addition, of the 53 duplicates classified as “Conserved” by CDROM, 18 are classified as “Neofunctionalized child” and 14 as “Specialized” by `CLOUD`. Finally, of the 41 duplicates classified as “Specialized” by `CDROM`, 18 are classified as “Neofunctionalzied child” by `CLOUD`. Taken together, these discrepancies reflect differences in the abilities of the `CLOUD` and `CDROM` classifiers to handle gene expression stochasticity.

**Figure 5:**
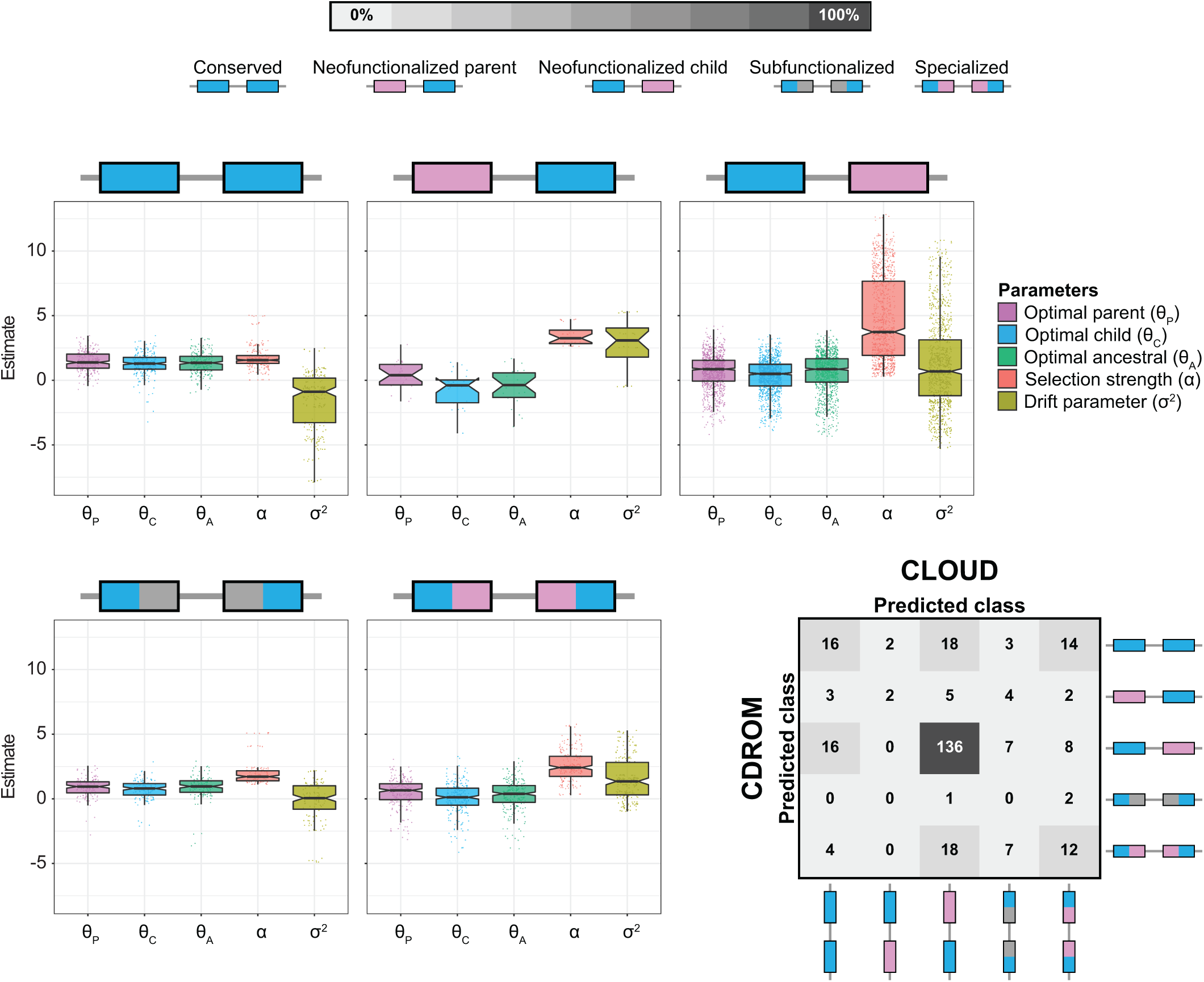
Classification and prediction results for application of `CLOUD` with *L* = 2 hidden layers to empirical data from *Drosophila* [Assis and Bachtrog, 2013, *Assis, 2019].* Box plots overlaid onto strip plots show distributions of log-transformed parameter estimates for each of the five classes of duplicate gene retention mechanisms. The confusion matrix in the bottom right illustrates the high concordance in classifications of `CLOUD` and `CDROM` for these empirical data, with both methods classifying the majority of duplicate genes as retained by neofunctionalization of the child copy.

We next examined the parameter predictions of `CLOUD`. Here, our major question was whether these predictions match theoretical expectations of duplicate gene retention mechanisms. To answer this question, we examined distributions of empirical parameter estimates for each class obtained with the `CLOUD` classifier (Figure 5). We first considered optimal expression estimates *θ*_P_, *θ*_C_, and *θ*_A_. For the “Conserved” class, the distributions of estimated *θ*_P_, *θ*_C_, and *θ*_A_ are not significantly different from one another, consistent with expectations (Table 2). For the “Neofunctionalized parent” class, the distribution of *θ*_P_ is elevated (though not significantly) relative to those of *θ*_C_ and *θ*_A_, whereas the distributions of *θ*_C_ and *θ*_A_ are not significantly different from one another. This qualitative pattern is as expected (Table 2), with the lack of a significant elevation of *θ*_P_ likely due to the small number of duplicate genes in this class. For the “Neofuntionalized child” class, the distribution of *θ*_C_ is significantly decreased relative to those of *θ*_P_ and *θ*_A_, whereas the distributions of *θ*_P_ and *θ*_C_ are not significantly different from one another, also consistent with theoretical expectations (Table 2). For the “Subfuntionalized” class, distributions of *θ*_P_ and *θ*_C_ are different (though not significantly) from one another, with the center of the distributions of *θ*_P_ and *θ*_C_ located at approximately the center of the *θ*_A_ distribution. This qualitative pattern matches expectations, though formal tests of significance were again underpowered due to only a handful of duplicates classified as “Subfunctionalized”. Finally, for the “Specialized” class, *θ*_P_, *θ*_C_, and *θ*_A_ are all significantly different from one another, matching theoretical expectations (Table 2). Analogously, we also observe a general concordance between estimates of *α* and *σ*^2^ and theoretical expectations of classified duplicate gene retention mechanisms. In particular, duplicate genes classified as neofunctionalized or specialized have significantly elevated estimated selection strengths (*α*) compared to those classified as conserved or subfunctionalized (Figure 5). These differences are consistent with theoretical expectations, as both neofunctionalization and specialization result in acquisitions of new functions that are hypothesized to be driven by strong selection, whereas both conservation and subfunctionalization result in preservations of ancestral functions that may occur in the absence of selection. Further, estimates of phenotypic drift (*σ*^2^) are also significantly larger for duplicate genes classified as neofunctionalized or specialized than as conserved or specialized. This result supports the hypothesis that traits require some minimum threshold of plasticity to effectively explore the space of novel phenotypes on which selection may act.

## Discussion

In this study, we have demonstrated that modeling of expression evolution and application of modern statistical learning techniques substantially enhances performance in classifying the retention mechanisms of duplicate genes and predicting their underlying parameters. Specifically, our new method `CLOUD` has high power and accuracy in discriminating among five classes of duplicate gene retention mechanisms (Figures 3, S4-S6), and high accuracy in parameter estimation (Figure 4). It represents a major advancement over the only previously available method, `CDROM` [Assis and Bachtrog, 2013, Perry and Assis, 2016], which has much lower classification power and accuracy (Figure 3), displays strong classification bias (Figures S3 and S4), and cannot perform the task of parameter prediction at all. Thus, `CLOUD` is currently the best method for classifying duplicate gene retention mechanisms from their expression data, and the only method for predicting the parameters driving their evolution. Moreover, though our study focuses on its application to gene expression data from multiple tissues, `CLOUD` can also be applied to gene expression data from multiple conditions of different types (*e.g.*, developmental stages or disease states), or even to gene expression data from a single condition, which is always the case for single-celled organisms. As a result, `CLOUD` can be used to learn about evolution after gene duplication in many diverse biological systems.

When designing the multi-layer neural network architecture of `CLOUD`, we took measures to mitigate overfitting through elastic net-style regularization [Zou and Hastie, 2005], which shrinks model weights through a mixture of *L*_1_- and *L*_2_-norm penalties [Hastie et al., 2009]. However, several other approaches, such as early stopping [Bishop, 1995, Sjöberg and Ljung, 1995, Goodfellow et al., 2016] and dropout [Srivastava et al., 2014, Goodfellow et al., 2016], could have been used instead to achieve a similar goal. Of the two alternatives mentioned, the dropout regularization procedure is closer to our approach, with a key difference in that regularization proceeds in a more stochastic fashion. Specifically, regularization is performed by dropping some proportion *x* ∈ (0, 1) of hidden units uniformly at random in each layer during each training epoch, thereby ensuring that fewer model parameters (weights) are trained during

each round of training. The optimal proportion *x* would then be chosen through cross-validation [Hastie et al., 2009], with all hidden units subsequently used during testing. Another option for reducing overfitting is ensembling [Breiman, 1994, Freund and Shapire, 1996a,b, Ridgeway, 1999] of neural networks (of which dropout is a specific form), either through bagging or boosting across neural networks [Schwenk and Bengio, 1998, Goodfellow et al., 2016], or by boosting across hidden layers of a neural network [Bengio et al., 2006]. Other ensemble approaches, such as random forests [Breiman, 2001, Hastie et al., 2009] and boosted regression and classification trees [Ridgeway, 1999, Hastie et al., 2009] may represent complementary flexible alternative frameworks to the neural network procedure employed here. In particular, they may be beneficial if expression data were absent for some tissues or genes, as they are able to naturally handle missing data [Hastie et al., 2009]. Though we considered these other regularization forms, we chose to utilize elastic net-style regularization as we felt that it provided a natural and deterministic mechanism for controlling model complexity.

We also considered an alternative approach for constructing the `CLOUD` classifier and predictor by employing maximum likelihood estimation [Casella and Berger, 2002, Brawand et al., 2011, Clavel et al., 2015]. Specifically, given expression data for parent, child, and ancestral genes, one can use maximum likelihood to estimate the set of parameters {*θ*_P_, *θ*_C_, *θ*_A_, *α, σ*^2^} from an OU model of expression evolution for each of the five retention mechanism classes, where optimal expression states (*θ*_P_, *θ*_C_, *θ*_A_) are constrained as shown in Table 2. Then, one can utilize likelihood ratio tests between models to derive a decision tree (similar to that used by `CDROM`) for performing classification. For these tests, the “Conserved” class would be nested within the “Neofunctionalized parent” and “Neofunctionalized child” classes, “Neofuntionalized parent” and “Neofunctionalized child” classes would be nested within the “Subfunctionalized” class, and the “Sub-functionalzied” class would be nested within the “Specialized” class. This procedure would require that model selection is performed using appropriate significance cutoffs [Casella and Berger, 2002], accounting for multiple testing [Neyman and Pearson, 1928]. Furthermore, the fit of the likelihood model would be highly dependent on underlying assumptions (*e.g.*, independence among tissues and gene tree estimates), and the “Specialized” model with five free parameters per tissue (Table 2) would be over-parameterized without including expression data from a fourth gene in an outgroup species. Additionally, it would be difficult to directly incorporate the genome-wide distribution of expression differences at single-copy genes to use as a baseline level of expression divergence. For these reasons, we believe that the framework implemented by `CLOUD` represents a more appropriate, powerful, and flexible approach for learning evolutionary retention mechanisms and parameters of duplicate genes.

Application of `CLOUD` to empirical data from *Drosophila* [Assis and Bachtrog, 2013, Assis, 2019] recapitulated many of the classifications previously inferred by `CDROM` [Assis and Bachtrog, 2013], notably classifying the majority of duplicate genes as retained by neofunctionalization in which the child copy acquires a new function (Figure 5). Predicted parameters of *Drosophila* duplicate genes were also generally concordant with theoretical expectations of their classified retention mechanisms (Table 2, Figure 5). In particular, observed differences among distributions of optimal expression estimates for parent (*θ*_P_), child (*θ*_C_), and ancestral (*θ*_A_) genes matched expectations for all retention mechanism classes (Table 2, Figure 5). Similarly, distributions of selection strength (*α*) estimates were shifted toward higher values for retention mechanisms in which there were acquisitions of new functions (neofunctionalization and specialization) relative to those in which ancestral functions were preserved (conservation and subfunctionalization, Figure 5), consistent with hypotheses that strong positive selection drives fixation of new functions after gene duplication [Ohno, 1970, He and Zhang, 2005, Rastogi and Liberles, 2005]. Interestingly, distributions of phenotypic drift (*σ*^2^) estimates were also elevated for classes in which there were acquisitions of new functions (Figure 5), perhaps because increased levels of phenotypic plasticity are necessary to explore new functions on which selection can act. This hypothesis is also supported by other studies of these *Drosophila* duplicate genes [Assis and Bachtrog, 2013], which found evidence of parallel sequence and expression evolution for all classified retention mechanisms [Assis and Bachtrog, 2013, Jiang and Assis, 2017]. Thus, our empirical findings are largely consistent both with long-held theoretical predictions [Ohno, 1970, He and Zhang, 2005, Rastogi and Liberles, 2005], and with results from previous analyses of these *Drosophila* duplicate genes [Assis and Bachtrog, 2013, Jiang and Assis, 2017]. Taken together, they illustrate that functional divergence after gene duplication in *Drosophila* is often asymmetric, tends to affect the child copy, and is driven by strong selection.

## Methods

In this section, we detail the algorithmic choices used to train `CLOUD`, the simulation setting used to compare its performance to the classifier `CDROM`, and the necessary steps for application of `CLOUD` and `CDROM` to empirical data from *Drosophila*.

### Training the neural network on data simulated from OU processes

Consider a set of *N*_*k*_ training observations for class *k* ∈ {1, 2, 3, 4, 5}, such that the total training sample size is *N* = *N*_1_ + *N*_2_ + *N*_3_ + *N*_4_ + *N*_5_. For observation *i* ∈ {1, 2, …, *N*} and output *k* ∈ {1, 2, …, *K*}, let 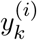 denote the true value and 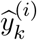 denote the estimated value. We wish to train a neural network model to minimize the overall discrepancy between 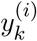 and 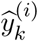 which we measure with the loss function 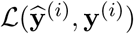, across the *N* samples and *K* outputs. Let

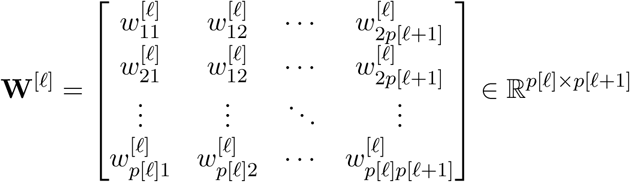

be the matrix of weights going from layer to layer + 1 for ∈ {0, 1, …, *L*}, and let

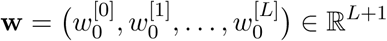

denote the vector of biases for all of the layers.

To train the neural network, we wish to identify the set of parameters (weights and biases) *𝒲* = {**w, W**^[0]^, …, **W**^[*L*]^} that minimize the cost

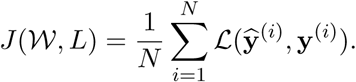

To prevent overfitting, we include an elastic-net style regularization penalty term [Zou and Hastie, 2005] on the weights of each layer with two tuning hyper parameters. Specifically, we reduce the complexity of the fitted model with the tuning parameter *λ* ≥ 0, which shrinks the magnitude of the weights to zero. We also perform simultaneous weight shrinkage and feature selection with the elastic net tuning parameter *γ* ∈ [0, 1], such that we are performing *L*_2_-norm regularization when *γ* = 0, *L*_1_-norm regularization that incorporates feature selection when *γ* = 1, and both types of regularization when *γ* ∈ (0, 1). In particular, we seek to find the model parameters *W* that minimize the penalized cost function

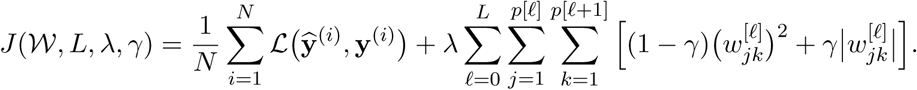

In the classification problem, for training observation *i* ∈ {1, 2, …, *N*}, we define the indicator random variable 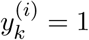 if observation *i* is from class *k*, and 0 otherwise. Hence, all output values are zero except for that corresponding to class *k*, which has a value of one. Based on this output, we employ the loss function used in *J* (*𝒲, L, λ, γ*) as the categorical cross entropy deviance [Goodfellow et al., 2016]

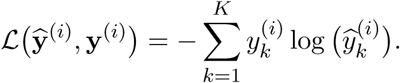

In the prediction problem, 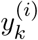 is instead the *k*th parameter value from simulated replicate *i*. Based on this output, we employ the loss function used in *J* (*𝒲, L, λ, γ*) as the residual sum of squared error

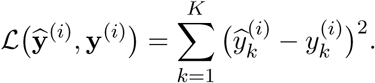

Simulated data have been successfully used to train models for learning about evolution from genomic data in many recent studies [Lin et al., 2011, Schrider and Kern, 2016, Sheehan and Song, 2016, Kern and Schrider, 2018, Schrider et al., 2018, Sugden et al., 2018, Flagel et al., 2019, Mughal et al., 2019, Mughal and DeGiorgio, 2019, Adrion et al., 2020]. Therefore, to train CLOUD for both classification and prediction, we generated a balanced simulated dataset with 10^4^ observations from each of the five classes, for a total of *N* = 50, 000 training observations. We assumed that tissues were independent, and that there were a total of *m* = 6 tissues as in an empirical data from *Drosophila* [Assis, 2019] that we later applied our method to, for a total of *p* = 108 input features.

To make the simulated dataset more realistic, we drew model parameters *T*_PC_ and *T*_PCA_ from empirical gene tree estimates for the set of *Drosophila* duplicate genes used by Assis and Bachtrog [2013]. The procedure for estimating these gene trees is detailed in subsection *Application of `CDROM` and `CLOUD` to empirical data from Drosophila* below. For all analyses, we scaled the root of the gene tree to have height one, and considered a new scaled time for the duplication event of *t*_PC_ = *T*_PC_*/T*_PCA_, such that *t*_PC_ represented the time of the duplication relative to the height of the root of the gene tree. For given class, we drew parameters 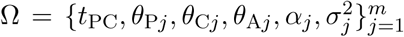 uniformly at random, assuming that *θ*_PC*j*_ = *θ*_PCA*j*_ = *θ*_A*j*_ for tissue *j* (schematic provided in Figure 2D). In particular, we drew *t*_PC_ from the distribution of empirical gene tree estimates, *θ*_*j*_ ∈ [−4, 4] for *j* ∈ {P, C, A}, *α* from log_10_(*α*) ∈ [0, 3], and *σ*^2^ from log_10_(*σ*^2^) ∈ [−2, 3]. We chose this specific range for *θ*_P_, *θ*_C_, and *θ*_A_ because we found that differences in log_10_-transformed empirical expression data were normally distributed, matching expectations under an OU model. For this reason, all empirical expression data were also log_10_-transformed prior to applying `CLOUD`. The class *k* is determined by {*θ*_P_, *θ*_C_, *θ*_A_}, which is summarized in Table 2. Then, we simulated gene expression data **e**^(*i*)^ ∈ ℝ^3*m*^ under model parameters for a given class *k* (Table 2), assuming independence among tissues, and generated *N*_*k*_ simulated replicates of parameter values 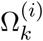 for *i* = 1, 2, …, *N*_*k*_.

Given a number of hidden layers *L* and the pair of regularization tuning parameters *λ* and *γ*, we estimated the set of parameters *𝒲* using the Adam optimizer [Kingma and Ba, 2014] with learning rate 10^−3^ and exponential decay rates for the first and second moment estimates of *β*_1_ = 0.9 and *β*_2_ = 0.999 [Kingma and Ba, 2014], respectively. This optimizer was used as it efficiently traverses the cost function surface *J* (*𝒲, L, λ, γ*) to rapidly identify the minimum [Kingma and Ba, 2014]. We also used mini-batch optimization with a batch size of 5000 observations for 500 epochs. To estimate *L, λ*, and *γ*, we performed five-fold cross-validation [Hastie et al., 2009]. Specifically, we used 80% (40,000) of observations for training, and held out the remaining 20% (10,000) for validation. We also ensured that the training and validation sets were balanced in class representation, such that there were equal numbers of observations from each class in the training (8,000) and validation (2,000) sets. To asses method performance for a given fold, we computed the validation loss

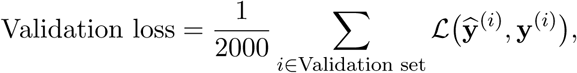

where the loss is either the categorical cross entropy deviance or the residual sum of squared error for the classifier or predictor, respectively [Goodfellow et al., 2016]. We then averaged this validation loss across the five folds to compute the cross-validation error [Hastie et al., 2009]. We considered values of *L* ∈ {0, 1, 2, 3} and *α* ∈ {0, 0, 0.1, …, 1.0}, as well as 25 values of *λ* chosen uniformly across log_10_(*λ*) ∈ [−12, −3]. Given the optimal cross-validation estimates 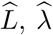, and 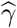 for *L, λ*, and *γ*, respectively, we estimated the neural network model parameters 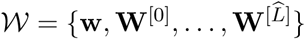 as

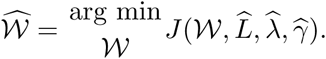

Previous studies have found that neural networks with enough hidden layers or units can approximate any function, and therefore lead to overfitting [Cybenko, 1989, Goodfellow et al., 2016]. Hence, based on simulations, we estimated that a neural network with 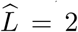 hidden layers provides the best cross-validation performance, with the validation loss for the classifier of approximately 0.918 with optimal tuning parameters 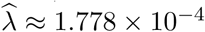 and 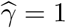, and the validation loss for the predictor of approximately 0.899 with optimal tuning parameters 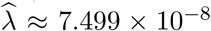 and 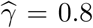. Comparisons of classification and prediction performances across the four network architectures *L* ∈ {0, 1, 2, 3} are given in Figures S1 and S2, highlighting the generally superior performance of the architecture with two hidden layers.

### Application of CDROM and CLOUD to simulated test data

To compare the relative classification powers and accuracies of CDROM and `CLOUD` and explore the prediction accuracy of `CLOUD`, we simulated training and test datasets for duplicate genes based on an OU process, which is described in subsection *Training the neural network on data simulated from OU processes* above. However, in that subsection, we assumed that expression vectors for single-copy genes *𝒢* were given. These would typically be extracted from the genome-wide distribution of single-copy genes for the pair of species being studied, such that trained models are based on the level of expression divergence typically observed in the study system.

For our simulated training and test sets, we generated a background set of 10,000 six-tissue expression vectors for single-copy genes that was inspired by those of the single-copy genes identified in *Drosophila* [Assis and Bachtrog, 2013, Assis, 2019]. Specifically, we applied the Brownian motion model [Felsenstein, 1973] implemented in mvmorph [Clavel et al., 2015] to expression vectors of single-copy genes between Species 1 and 2, assuming that tissues were independent and that the root of the two-species phylogeny had height one, to estimate ancestral expression *θ* and variance *σ*^2^ parameters consistent with the empirical distribution in *Drosophila* at each tissue and single-copy gene. Given the set of parameters, we then sampled values of *θ* and *σ*^2^ uniformly at random from the estimated empirical distribution, and generated simulated single-copy expression vectors in Species 1 and Species 2 for *m* = 6 independent tissues, giving us the simulated set *𝒢*.

To test either the classifier or predictor, we generated a balanced set of duplicate gene expression vectors, such that each of the five classes had 1, 000 observations, for a total of *N* = 5, 000 test observations. We assumed that tissues were independent, and that there was a total of *m* = 6 tissues as in the training set. For a given class, we drew parameters 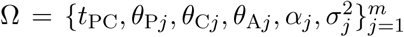 uniformly at random. In particular, as with the training set, we drew *t*_PC_ from the distribution of empirical gene tree estimates, *θ*_*j*_ ∈ [−4, 4] for *j* ∈ {P, C, A}, *α* from log_10_(*α*) ∈ [0, 3], and *σ*^2^ from log_10_(*σ*^2^) ∈ [−2, 3]. The class *k* was determined by {*θ*_P_, *θ*_C_, *θ*_A_} (Table 2), and gene expression data were generated **e**^(*i*)^ ∈ ℝ^3*m*^ under model parameters for a given class *k* (Table 2), assuming independence among tissues and *N*_*k*_ simulated replicates of parameter values 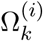 for *i* = 1, 2, …, *N*_*k*_.

To assay how `CLOUD` performs in different portions of the parameter space, we also examined its accuracy on test sets drawn from restricted parameter values. Specifically we considered three distinct ranges for *α* of [1, 10], [10, 100], and [100, 1000], and five distinct ranges for *σ*^2^ of [0.01, 0.1], [0.1, 1], [1, 10], [10, 100], and [100, 1000]. For each combination of a range for *α* and a range for *σ*^2^, we sampled *α* and *σ*^2^ uniformly at random.

### Application of CDROM and CLOUD to empirical data from *Drosophila*

We applied `CDROM` and `CLOUD` to empirical data consisting of *Drosophila* duplicate and single-copy genes [Assis and Bachtrog, 2013] and their expression abundances in six tissues [Assis, 2019]. Because `CLOUD` requires estimates of *T*_PC_ and *T*_PCA_, we first generated multiple sequence alignments with MACSE [Ranwez et al., 2018], which accounts for underlying codon structure, and then inferred a gene tree with PhyML [Guindon et al., 2010] for each parent, child, and ancestral gene in the duplication dataset [Assis and Bachtrog, 2013]. The empirical distributions of estimated *T*_PC_ and *T*_PCA_ across these gene trees were used as input to an OU process to generate a balanced training set with *N* = 50, 000 observations as described in subsection *Training the neural network on data simulated from OU processes* above. Gene expression data from single-copy genes in *Drosophila* [Assis and Bachtrog, 2013, Assis, 2019] were used as the set *G* necessary for application of both `CDROM` and `CLOUD`. We trained a classifier and predictor for `CLOUD` assuming *L* = 2 hidden layers, and through five-fold cross validation [Hastie et al., 2009], estimating the regularization tuning parameters as 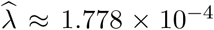 and 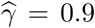 for the classifier and 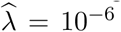 and 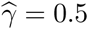 for the predictor.

## Acknowledgments

This work was supported by National Science Foundation grants DEB-1555981 and DEB-2001059 to RA and DEB-1949268 and BCS-2001063 to MD, and by National Institutes of Health grant R35-GM128590 to MD.

