## Supplementary Materials for "Learning retention mechanisms and evolutionary parameters of duplicate genes from their expression data"

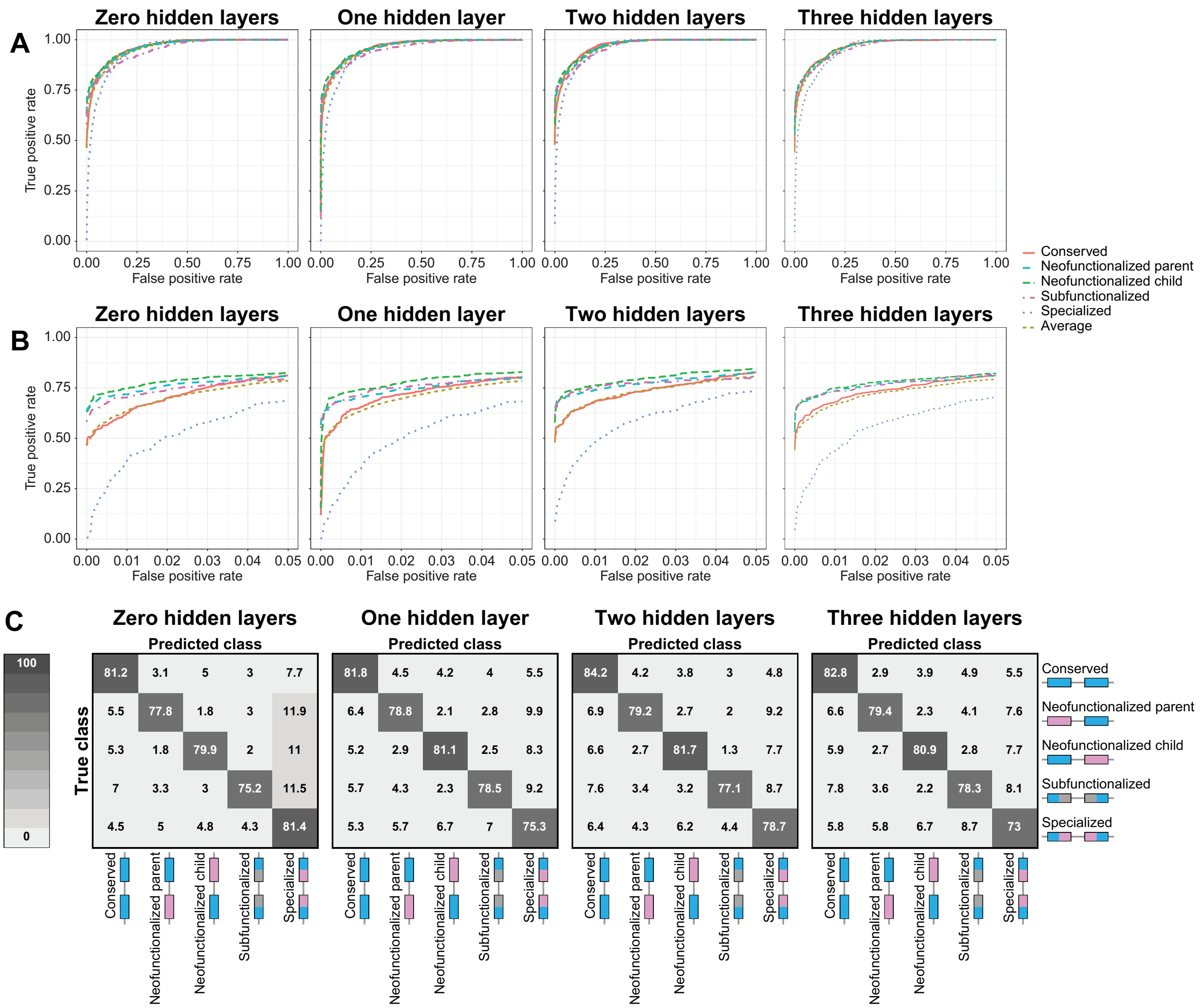


Figure S1: Classification results for applications of CLOUD with $L \in \{0, 1, 2, 3\}$ hidden layers to data simulated under parameters $\alpha \in [1, 10^3]$ and $\sigma^2 \in [10^{-2}, 10^3]$. (A) Receiver operating characteristic curves across the full range of false positive rates. (B) Receiver operating characteristic curves truncated at a false positive rate of 5%. (C) Confusion matrices depicting classification rates.

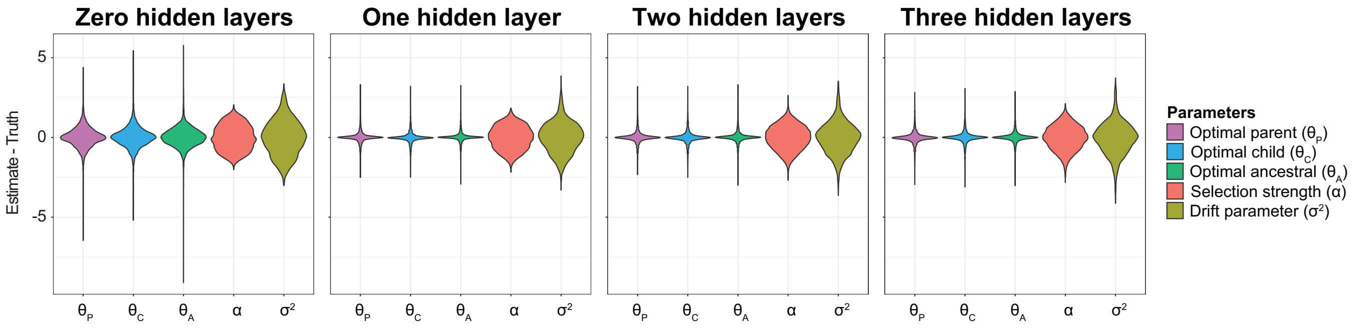


Figure S2: Prediction results for applications of `CLOUD` with $L \in \{0, 1, 2, 3\}$ hidden layers to data simulated under parameters $\alpha \in [1, 10^3]$ and $\sigma^2 \in [10^{-2}, 10^3]$. Violin plots display distributions of parameter prediction errors averaged across the $m = 6$ tissues for each simulated test set.

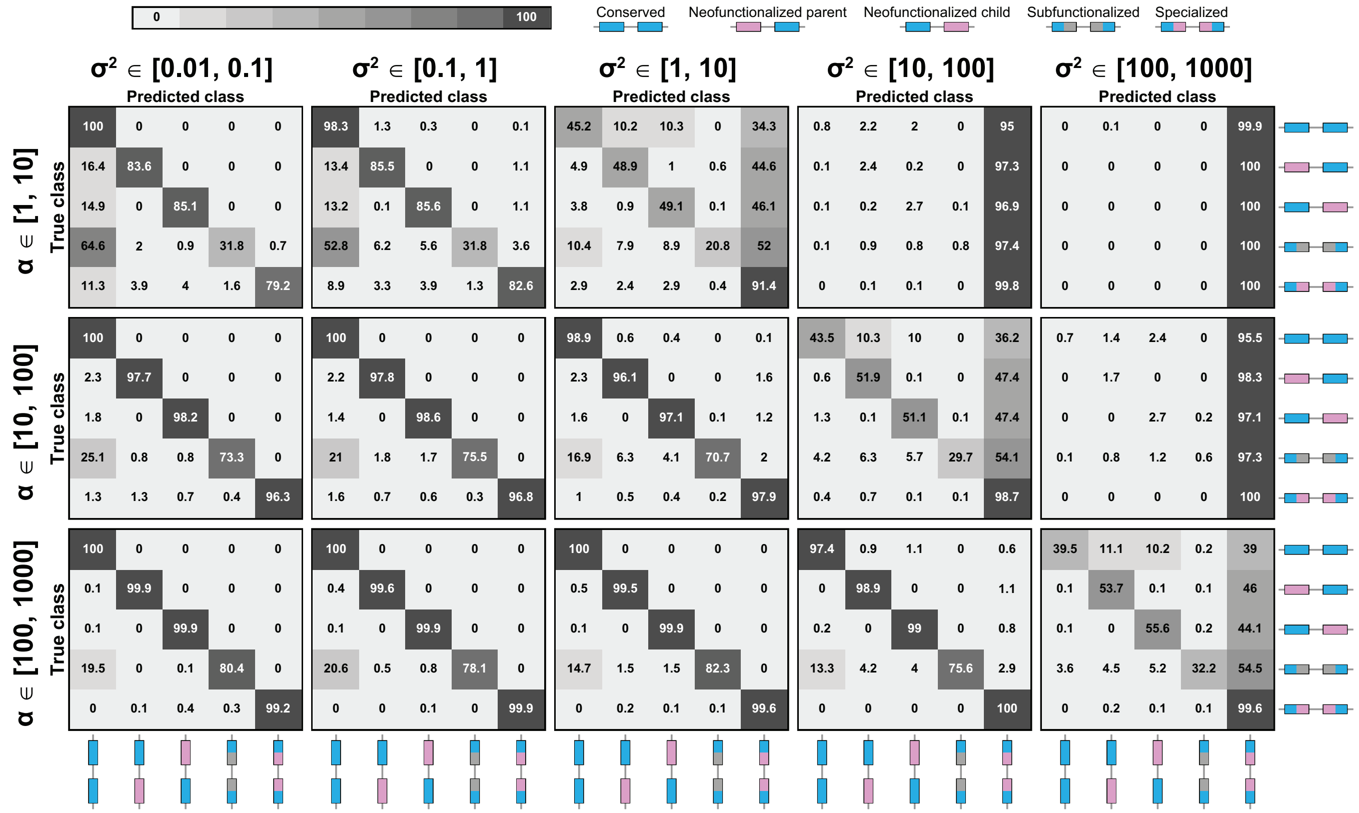


Figure S3: Confusion matrices for applications of `CDROM` to data simulated under specific parameter ranges for $\alpha$ and $\sigma^2$. Classification accuracy is highest for large $\alpha$ and small $\sigma^2$, and lowest for small $\alpha$ and large $\sigma^2$.

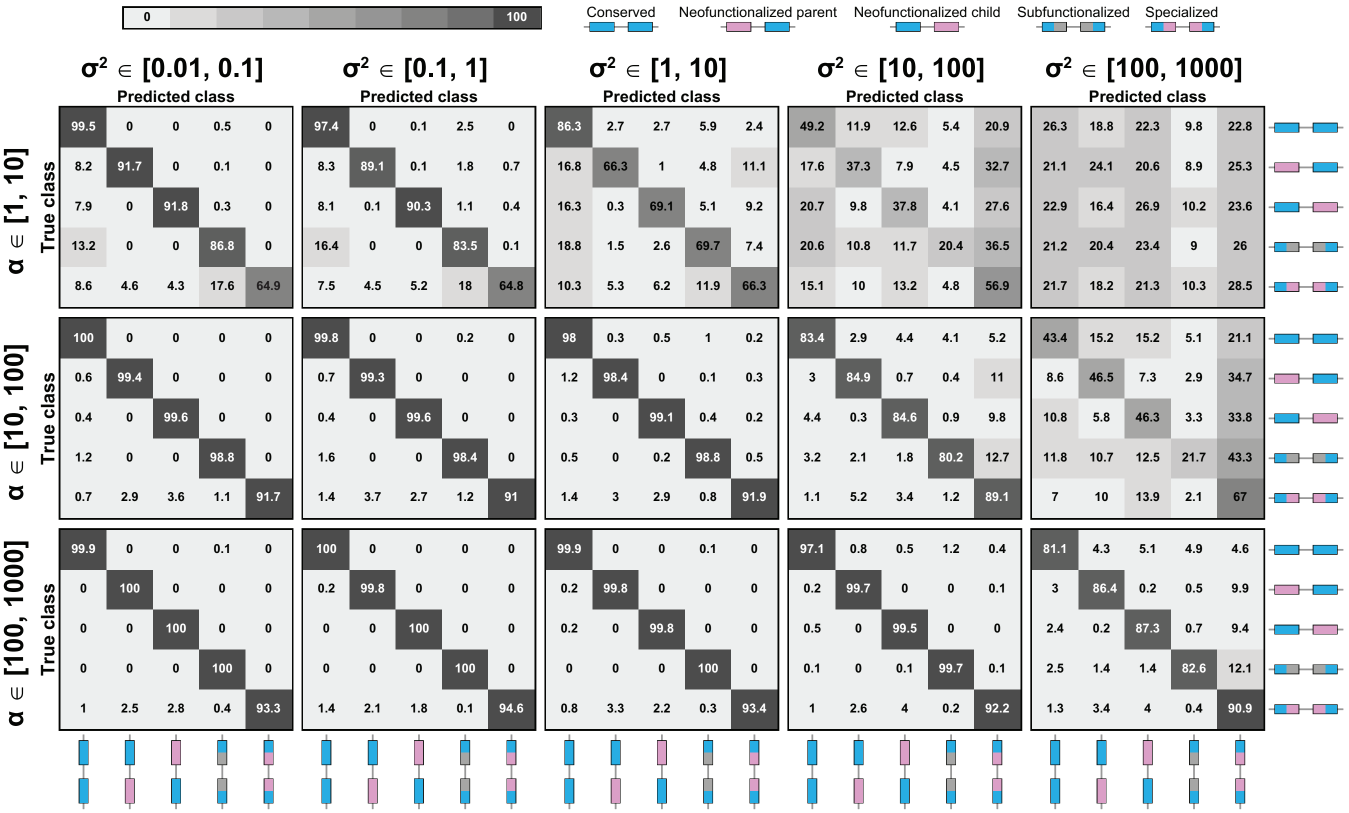


Figure S4: Confusion matrices for applications of CLOUD with $L = 2$ hidden layers to data simulated under specific parameter ranges for $\alpha$ and $\sigma^2$. Classification accuracy is highest for large $\alpha$ and small $\sigma^2$, and and lowest for small $\alpha$ and large $\sigma^2$.

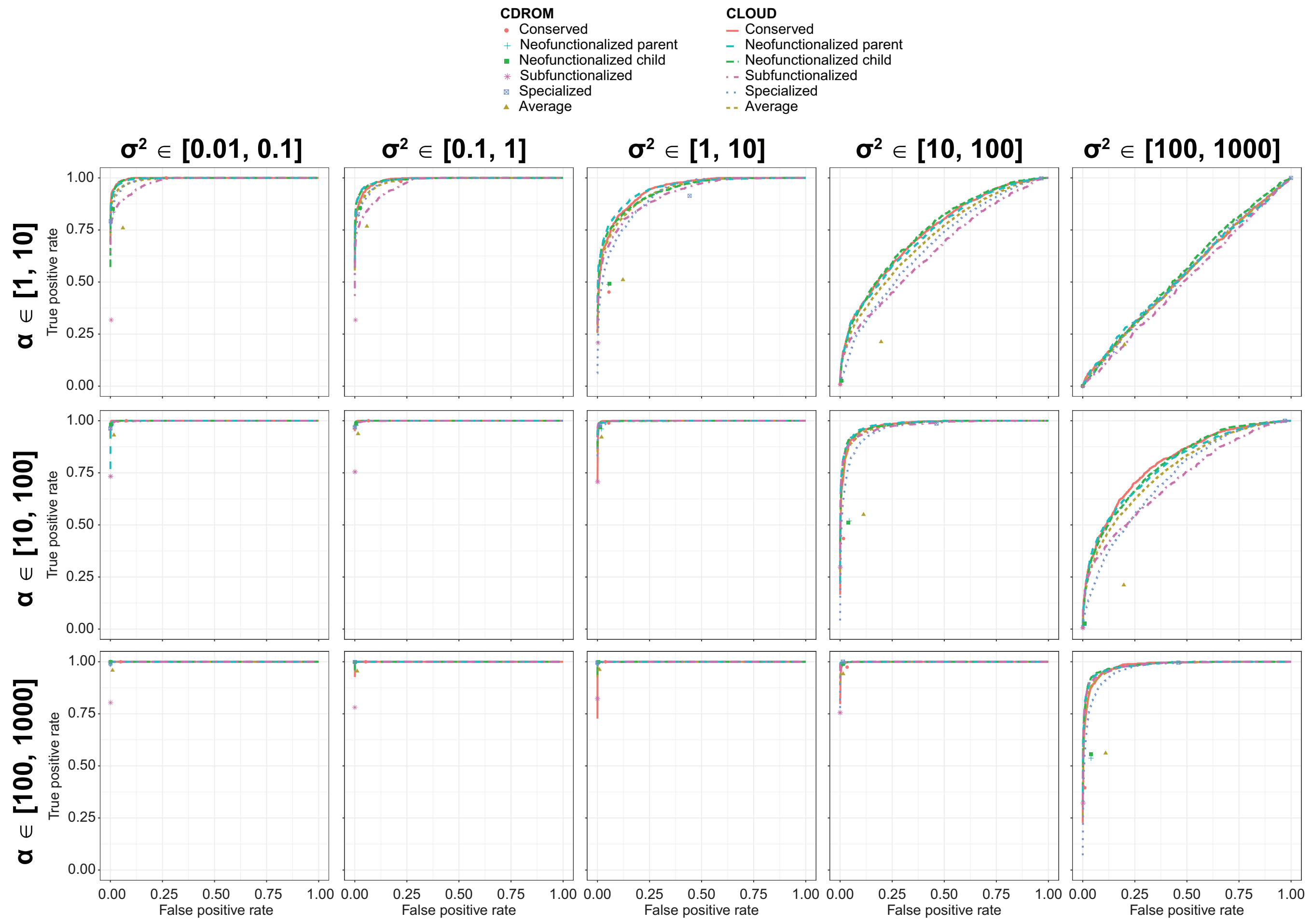


Figure S5: Receiver operating characteristic curves for applications of `CDROM` and `CLOUD` with $L = 2$ hidden layers to data simulated under specific parameter ranges for $\alpha$ and $\sigma^2$. Classification accuracy is highest for large $\alpha$ and small $\sigma^2$, and lowest for small $\alpha$ and large $\sigma^2$. Because `CDROM` is a decision tree classifier, its true positive and false positive rates are plotted as points.

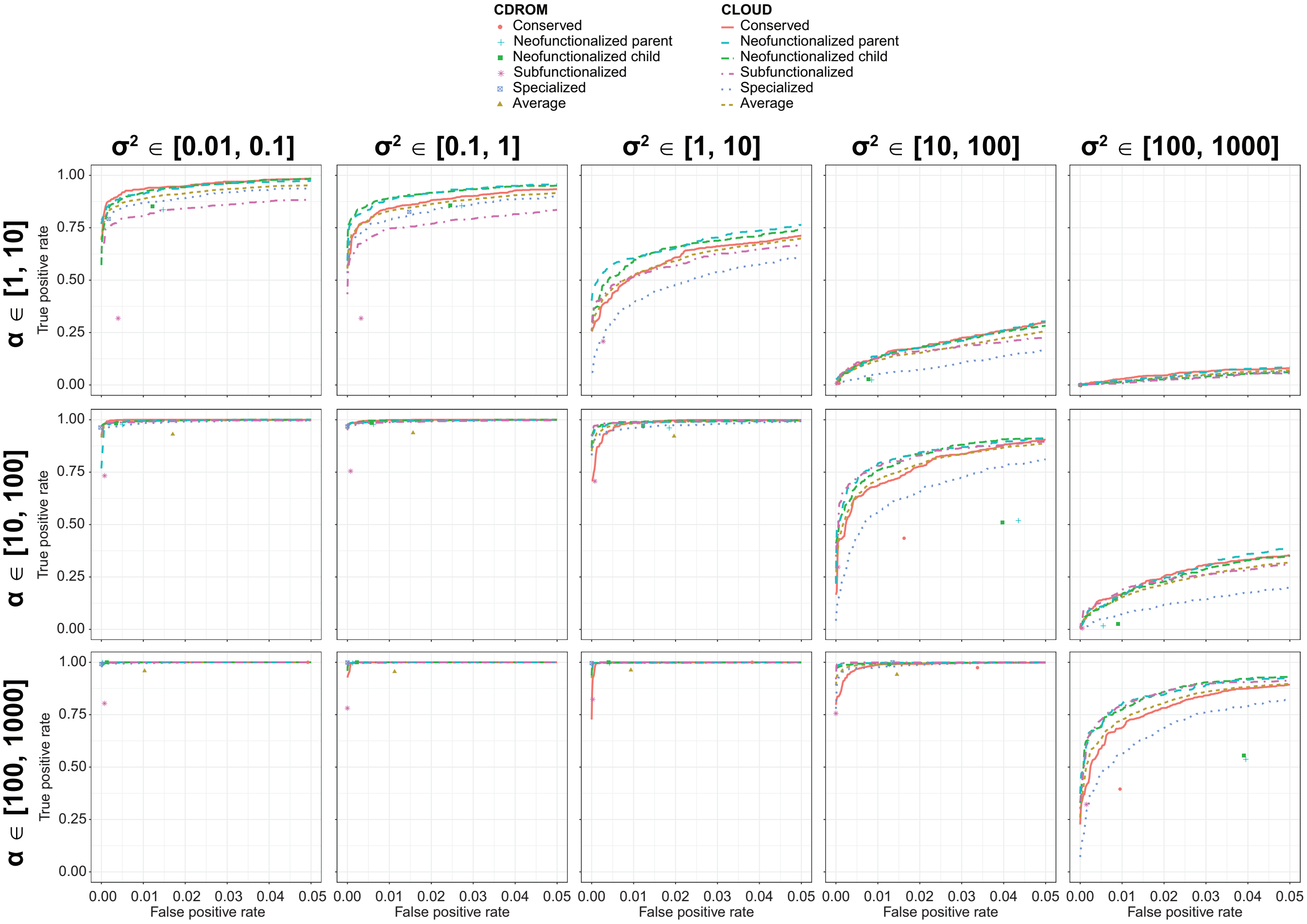


Figure S6: Receiver operating characteristic curves truncated at a false positive rate of 5% for applications of CDROM and CLOUD with $L = 2$ hidden layers to data simulated under specific parameter ranges for $\alpha$ and $\sigma^2$. Classification accuracy is highest for large $\alpha$ and small $\sigma^2$, and and lowest for small $\alpha$ and large $\sigma^2$. Because CDROM is a decision tree classifier, its true positive and false positive rates are plotted as points.

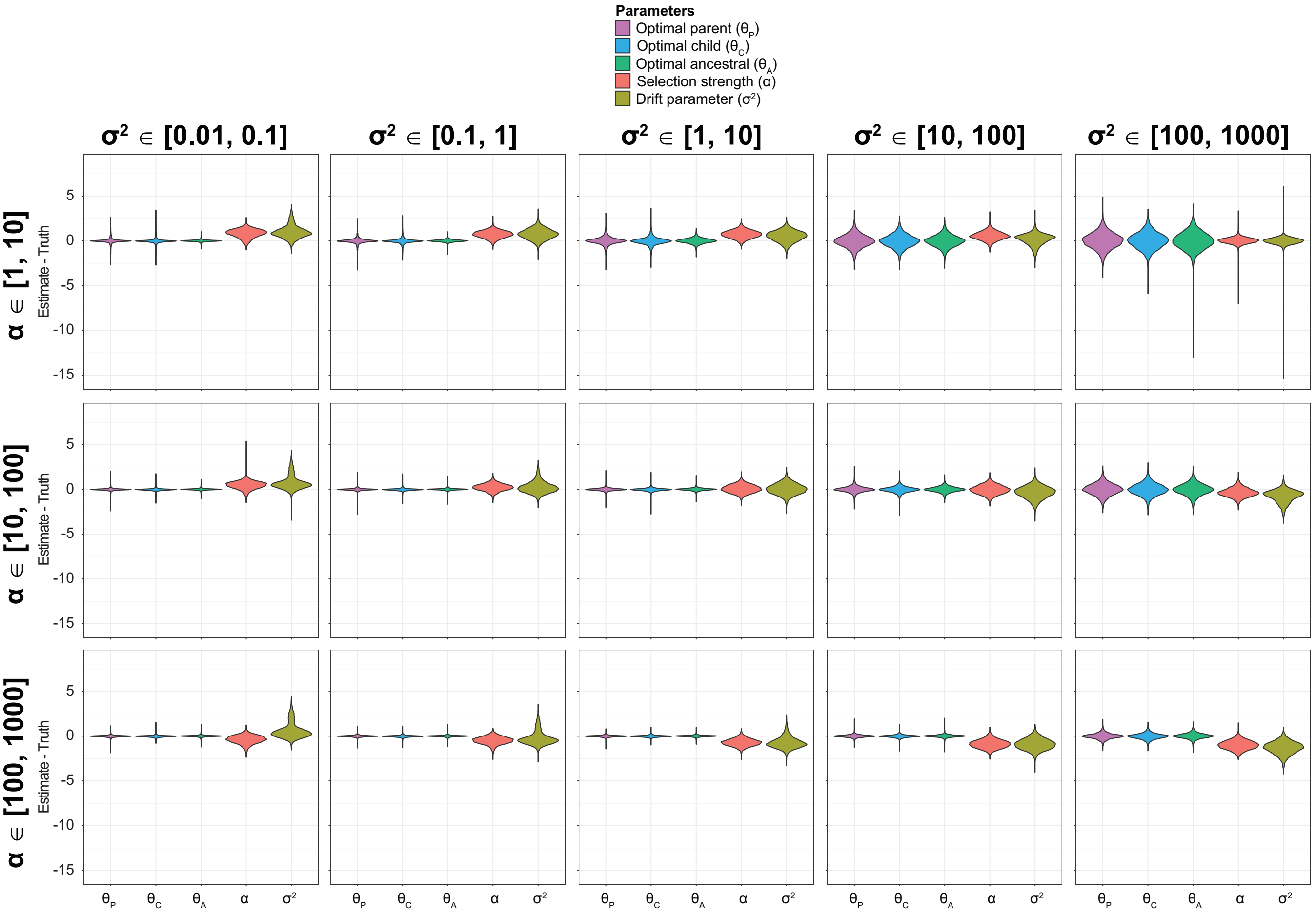


Figure S7: Prediction results for applications of `CLOUD` with $L = 2$ hidden layers to data simulated under specific parameter ranges for $\alpha$ and $\sigma^2$. Violin plots display distributions of parameter prediction errors averaged across across the $m = 6$ tissues for each simulated test set.
